# Longitudinal Study of Leukocyte DNA Methylation and Biomarkers for Cancer Risk in Older Adults

**DOI:** 10.1101/597666

**Authors:** Alexandra H. Bartlett, Jane W Liang, Jose Vladimir Sandoval-Sierra, Jay H Fowke, Eleanor M Simonsick, Karen C Johnson, Khyobeni Mozhui

## Abstract

**Background:** Changes in DNA methylation over the course of life may provide an indicator of risk for cancer. We explored longitudinal changes in CpG methylation from blood leukocytes, and likelihood of a future cancer diagnosis.

**Methods:** Peripheral blood samples were obtained at baseline and at follow-up visit from 20 participants in the Health, Aging and Body Composition prospective cohort study. Genome-wide CpG methylation was assayed using the Illumina Infinium Human MethylationEPIC (HM850K) microarray.

**Results:** Global patterns in DNA methylation from CpG-based analyses showed extensive changes in cell composition over time in participants who developed cancer. By visit year 6, the proportion of CD8+ T-cells decreased (p-value = 0.02), while granulocytes cell levels increased (p-value = 0.04) among participants diagnosed with cancer compared to those who remained cancer-free (cancer-free vs. cancer-present: 0.03 ± 0.02 vs. 0.003 ± 0.005 for CD8+ T-cells; 0.52 ± 0.14 vs. 0.66 ± 0.09 for granulocytes). Epigenome-wide analysis identified three CpGs with suggestive p-values ≤ 10^−5^ for differential methylation between cancer-free and cancer-present groups, including a CpG located in *MTA3*, a gene linked with metastasis. At a lenient statistical threshold (p-value ≤ 3 × 10^−5^), the top 10 cancer-associated CpGs included a site near *RPTOR* that is involved in the mTOR pathway, and the candidate tumor suppressor genes *REC8, KCNQ1*, and *ZSWIM5*. However, only the CpG in *RPTOR* (cg08129331) was replicated in an independent data set. Analysis of within-individual change from baseline to Year 6 found significant correlations between the rates of change in methylation in *RPTOR*, *REC8* and *ZSWIM5*, and time to cancer diagnosis.

**Conclusion:** The results show that changes in cellular composition explains much of the cross-sectional and longitudinal variation in CpG methylation. Additionally, differential methylation and longitudinal dynamics at specific CpGs could provide powerful indicators of cancer development and/or progression. In particular, we highlight CpG methylation in the *RPTOR* gene as a potential biomarker of cancer that awaits further validation.

## Background

DNA methylation plays a central role in cell differentiation and in defining cellular phenotypes. Differences in DNA methylation have been associated with a growing list of morbidities, ranging from metabolic disorders and age-related decline in health, to developmental and neuropsychiatric conditions. The standard approach in an epigenome-wide association study (EWAS), which attempts to link DNA methylation to disease, involves collection of a single biospecimen from each participant (typically peripheral blood or saliva) and performing cross-sectional analyses to compare methylation patterns in cases against matched healthy controls [1, 2]. While differences in CpG methylation between cases and controls may be directly related to disease, these case-control differences may also represent DNA sequence variation, differences in disease treatment, differences in behavior or environment, or differences in cellular composition [3, 4]. Despite these limitations in the interpretation of DNA methylation results, such epigenetic markers, if consistent and replicable, could serve as powerful biomarkers that can be assayed from minimally invasive tissues such as circulating blood.

Cancer is fundamentally due to abnormal cell phenotype and proliferation, and historically, it was the first disease linked to aberrant DNA methylation [5–7]. The cancer epigenome often involves global hypomethylation at repetitive elements, while also potentially involving the hypermethylation at CpGs in the promoter regions of tumor suppressor genes and other cancer-related genes [8–10]. While abnormal epigenomic changes within tumor cells would hold the most impact, there is developing evidence that methylation changes relevant to cancer progression can be detected in circulating blood. For example, global changes in repetitive elements as well as targeted CpG methylation found in DNA from blood cells have been reported for multiple cancer types [11–15]. This suggests the possibility of a pan-cancer biomarker panel detectable in blood that could precede the clinical detection and diagnosis of cancer [16].

Few longitudinal studies have investigated the time-dependent dynamics in DNA methylation as a potentially important indicator of tumorigenesis [14, 15]. The present study examines the longitudinal restructuring of the methylome over five years and evaluates whether change in CpG methylation is a biomarker of cancer in older adults. Our approach involves dimension reduction techniques and evaluates leukocyte proportions and differential methylation at the level of individual CpGs. Overall, our study defined global and targeted changes in the blood methylome that were correlated to cellular composition, aging, and cancer in the Health ABC cohort.

## Methods

### Health, Aging and Body Composition Study (Health ABC Study)

The Health ABC Study is a prospective, longitudinal cohort that was recruited in 1997–1998 and consisted of 3,075 older men and women participants aged 70– 79 years at baseline. Participants resided in either the Memphis, TN or Pittsburgh, PA metropolitan areas, and were either of African American or Caucasian ancestry [17]. Individuals with limited mobility, history of active treatment for cancer in the past 3 years, or with known life-threatening disease were excluded. More information on participant screening and recruitment can be found at the study website [18]. There were annual clinical visits to record health and function, and subjects were followed for up to 16 years. The study collected data on adjudicated health events, including cancer, and a biorepository was developed. All participants provided written informed consent and all sites received IRB approval. The present study leverages data on a small set of Health ABC participants who had DNA available from buffy coat collected at baseline and at follow-up visits (mostly at year 6 from baseline).

### DNA methylation microarray and data processing

Due to low DNA quality/quantity, 3 participants had DNA from only one visit year, and in total, we generated DNA methylation data on 37 samples. Participant characteristics and DNA collection time-points are provided in **Table 1**. Seven of the 20 participants received adjudicated cancer diagnosis in following years with four between baseline and Year 6, and three after Year 6.

**Table 1.** Characteristics of participants.

| ID | Ancestry<sup>1</sup> | Sex<sup>1</sup> | Age<sup>1</sup> | Followup<br>Year<sup>2</sup> | Cancer<sup>3</sup> | Time<sup>4</sup> |
| --- | --- | --- | --- | --- | --- | --- |
| Per1 | EA | Male | 75 | 6 | no |  |
| Per2 | AA | Male | 71 | 6 | yes<sup>p</sup> | 7 |
| Per3 | AA | Female | 72 | 6 | no |  |
| Per4 | EA | Male | 74 | 6 | yes<sup>c</sup> | 5 |
| Per5 | EA | Female | 76 | 6 | no |  |
| Per6 | EA | Male | 75 | 6 | yes<sup>p</sup> | 4 |
| Per7 | AA | Male | 76 | 2 | no |  |
| Per8 | AA | Female | 78 | 6 | no |  |
| Per9 | EA | Female | 78 | 6 | yes<sup>b</sup> | 1 |
| Per10 | AA | Male | 74 | 6 | yes<sup>o</sup> | 10 |
| Per11 | AA | Female | 74 | 6 | no |  |
| Per12 | AA | Female | 71 | 6 | no |  |
| Per13 | EA | Male | 76 | 6 | yes<sup>l</sup> | 0.5 |
| Per14 | EA | Male | 75 | na | no |  |
| Per15 | EA | Female | 73 | 6 | no |  |
| Per16 | EA | Male | 73 | 6 | no |  |
| Per17 | AA | Female | 76 | na | no |  |
| Per18 | AA | Female | 78 | 6 | yes<sup>s</sup> | 11 |
| Per19 | AA | Female | 72 | 6 (no<br>baseline<br>DNA) | no |  |
| Per20 | EA | Female | 70 | 6 | no |  |

<sup>1</sup>Self-reported race, sex, and age at baseline; EA = European Americans or Caucasians and AA = African Americans
<sup>2</sup>Year from baseline when second DNA sample was collected; two participants had no follow-up DNA and one participant had no baseline (visit year 1) DNA due to low DNA quality/quantity. <sup>3</sup>Cancer diagnosis during following years; all participants were considered free of diagnosed cancer at time of screening and recruitment; p = prostate, c = colon; b = breast; l = leukemia; s = stomach; o = other <sup>4</sup>Time from baseline to cancer diagnosis in years

DNA methylation assays were performed, as per the manufacturer’s standard protocol, using the Illumina Infinium Human MethylationEPIC BeadChips (HM850K) (http://www.illumina.com/). For this work, samples were shipped to the Genomic Services Lab at the HudsonAlpha Institute for Biotechnology (http://hudsonalpha.org). The HM850K arrays come in an 8-samples-per-array format; prior to hybridization, samples were randomized so that individuals were randomly distributed across the arrays. Raw intensity data (idat files) were loaded to the R package, minfi (version 1.22) [19]. Methylation level at each CpG was estimated by the β-value, which is the ratio of fluorescent intensities between the methylated probe and unmethylated probe. For quality checks (QC), we compared the log median intensities between the methylated (M) and unmethylated (U) channels using the “plotQC” function and examined the density plots for the β-values (QC plots are provided in **Additional file 1: Figure S1**). All 37 samples passed the initial QC (**Additional file 1: Figure S1A)**. Participant sex, as determined by DNA methylation, matched the sex listed in the participant record.

Methylation data was quantile-normalized using the minfi “preprocessQuantile” function. To evaluate sample clustering, we performed hierarchical cluster analysis and principal component analysis (PCA) using the full set of 866,836 probes (**Additional file 1: Figure S1B**). Sex was a strong source of variance when the full set of probes was used. We therefore filtered out 19,681 probes that targeted CpGs on the sex chromosomes. An additional 2,558 probes were filtered out due to detection p-values > 0.01 in 3 or more samples. Finally, we excluded 104,949 probes that have been flagged as unreliable due to poor mapping quality or overlap with genetic sequence variants (MASK.general list of probes from [20]). This resulted in 739,648 probes that were considered for downstream analyses. The updated PC plot showed no clustering by sex or by the Illumina Sentrix ID, which indicated that there was no strong chip effect. However, there were two outlier samples from the same individual (Per13) (**Additional file 1: Figures S1B, S1C**). Since the two samples were assayed on different Sentrix arrays, the outlier status is unlikely to be the result of technical artifact, but rather, flags Per13 as a biological outlier (excluded from downstream analyses). As an additional error checking step to confirm if samples from the same participants paired appropriately with self, we repeated the unsupervised cluster analysis using only 52,033 probes that were filtered out from the main set of probes due to overlap with common single nucleotide polymorphism (SNP) in the dbSNP database (**Additional file 2: Figures S2**).

### Estimating cellular composition

Cellular heterogeneity has a strong influence on DNA methylation, and methods have already been developed to estimate cellular composition of whole blood from genome-wide DNA methylation data [21–23]. We used the “estimateCellCounts” function in minfi, which implements a modified version of the algorithm by Houseman et al. [23] and relies on a panel of cell-type specific CpGs to serve as proxies for different types of white blood cells.

### Analyses of DNA methylation data

Considering the small sample size of the genome-wide data, we first started with a dimension reduction approach and applied PCA to capture the major sources of global variance in the methylome. The top 5 principal components (PCs) were then related to baseline variables using chi-squared tests for categorical variables (sex and race), and analysis of variance for continuous variables (BMI and age). We also examined the time-dependent change in the PCs with visit year as the predictor variable. Correlations between leukocyte types and the PCs were examined using bivariate analysis. We considered adjudicated cancer diagnosis as the main outcome variable and examined whether methylome-based variables differed between those who developed cancer and those who remained cancer-free.

Our primary analysis was to evaluate differential methylation at the CpG-level. As in Roos et al. [16], we first fitted a linear regression model on each probe for the first 5 PCs (β-value ∼ PC1 + PC2 + PC3 + PC4 + PC5) to adjust for the effects of confounding variables such as cellular heterogeneity and additional unknown sources of variance. The adjusted β-values were then used to examine differential methylation between cancer-free and cancer-present groups using t-tests. The t-tests were done with data only from visit Year 6. To evaluate the reliability of identified cancer-associated CpGs, we acquired the full results from Roos et al. [16], and compared the p-values and the direction of effect (i.e., increases or decreases in methylation in the cancer group relative to cancer-free group). To evaluate longitudinal trajectory, we considered only the top 10 CpGs associated with cancer and calculated the change in β-values from baseline to Year 6 (deltaβ = Year 6 – baseline), which was then correlated to time-to-diagnosis (i.e., years from baseline to when participant received diagnosis).

### Data availability

The deidentified raw data set with normalized β-values and EWAS statistics will be deposited to the NCBI NIH Gene Expression Omnibus (this will be made available upon acceptance by a peer-reviewed journal).

## Results

### Participant characteristics

The study sample included almost equal numbers of men and women, and equal numbers of African American and Caucasian participants (**Table 1**). Baseline age ranged from 70 to 78 years with an average age of 74 ± 2.4 years. Follow-up DNA collection occurred at Year 6, with the exception of one participant with follow up DNA collected at year 2 (Per7). Three participants had DNA from only one time point, and thus these were included in the cross-sectional analysis but not the time-dependent analysis.

During the Health ABC follow-up period, 7 participants (35%) were diagnosed with cancer at times ranging from 6 months to 11 years from baseline (**Table 1**). Cancer diagnoses included cancer of the prostate, colon, breast, and stomach, as well as one case of leukemia. There were no differences in race, sex, or baseline age or body mass index (BMI) between participants diagnosed with cancer and those who remained cancer-free (**Table 2**).

**Table 2.** Baseline characteristics of participants by cancer diagnosis.

|  | Cancer<sup>1</sup> |  | p-value<sup>2</sup> |
| --- | --- | --- | --- |
|  | no | yes |  |
| N | 13 (65%) | 7 (35%) |  |
| Age | 74 (±2.3) | 75 (±2.5) | 0.29 |
| Ancestry/race<sup>3</sup> |  |  | 0.64 |
| AA | 7 (35%) | 3 (15%) |  |
| EA | 6 (30%) | 4 (20%) |  |
| Sex |  |  | 0.08 |
| Female | 9 (45%) | 2 (10%) |  |
| Male | 4 (20%) | 5 (25%) |  |
| BMI | 27.01 (±3.77) | 27.75 (±5.42) | 0.72 |

<sup>1</sup>Counts (percent of total) for categorical variables and mean (SD) for continuous variables
<sup>2</sup>P-values based on Chi-square test and ANOVA
<sup>3</sup>Self-reported race identity; EA = European Americans or Caucasians, and AA = African Americans

### Quality of DNA methylation data and outlier identification

Unsupervised hierarchical clustering using the full set of probes showed that 15 of the individuals with longitudinal data paired within the same participant (**Additional file 1: Figure S1B**). The two exceptions, Per1 (cancer-free) and Per9 (received cancer diagnosis at year 1 from baseline), did not cluster with self, and this observation suggests potential intra-individual discordance in the epigenetic data or increased cellular heterogeneity over time [24, 25]. To verify that the non-pairing longitudinal samples are indeed from the same respective participants, we performed the cluster analysis using only probes that were flagged for overlap with SNPs, as these provide a signal for underlying genotype variation. Using these SNP probes, all individuals with longitudinal samples, including Per1 and Per9, paired appropriately with self (**Additional file 2: Figures S2**). Overall, the PC and cluster plots showed no batched effects and a generally stable methylation pattern over time, with the exception of the two participants. The QC analyses also identified Per13 as an outlier (**Additional file 1: Figures S1B, S1C**). Since Per13 was diagnosed with leukemia within 6 months of the first Health ABC visit, the distinct methylation pattern is consistent with disease-related changes in leukocyte composition, and Per13 was excluded from further analyses.

### Longitudinal changes in CpG-based blood cell composition

We performed a CpG-based estimation of blood cell proportions [21–23]. We evaluated differences in blood composition between baseline and Year 6. The estimated proportion of CD8+ T-cells decreased, while the proportion of granulocytes increased (**Figure 1A, 1B; Table 3**). The proportions of the other blood leukocyte subtypes remained relatively stable with no significant differences between the two visits (estimates for all participants at both time points are in **Additional file 3: Table S1**). We however note pronounced changes in cell composition for Per1, one of the two participants that did not pair with self in the hierarchical cluster; cellular heterogeneity partly explains the discordance in the longitudinal data.

**Table 3.** Association between cancer and CpG-based estimates of blood cells and PC1.

| Comparison between baseline and year 6<sup>†</sup> |  |  |  |  |  |  |
| --- | --- | --- | --- | --- | --- | --- |
|  | Baseline |  |  | Year 6 |  | p<br>(baseline vs 6) |
| CD8T | 0.07 ± 0.06 |  |  | 0.02 ± 0.02 |  | 0.006 |
| Gran | 0.46 ± 0.14 |  |  | 0.57 ± 0.14 |  | 0.02 |
| PC1 | -7.31 ± 15.7 |  |  | 8.51 ± 13.41 |  | 0.004 |
|  | Baseline (cancer yes vs. no) |  |  | Year 6 (cancer yes vs. no) |  |  |
| Cancer | No | Yes | p | No | Yes | p |
| CD8T | 0.08 ± 0.07 | 0.04 ± 0.02 | 0.16 | 0.03 ± 0.02 | 0.003 ± 0.005 | 0.02 |
| Gran | 0.43 ± 0.14 | 0.52 ± 0.12 | 0.17 | 0.52 ± 0.14 | 0.66 ± 0.09 | 0.04 |
| PC1 | -12.19 ± 13.73 | 2.45 ± 15.87 | 0.06 | 2.13 ± 10.99 | 19.14 ± 10.23 | 0.008 |

<sup>†</sup>Excludes Person 13 and data from Year 2
CD8T: CD8+ T-cells; Gran: granulocytes; PC1: principal component 1

**Figure 1.**
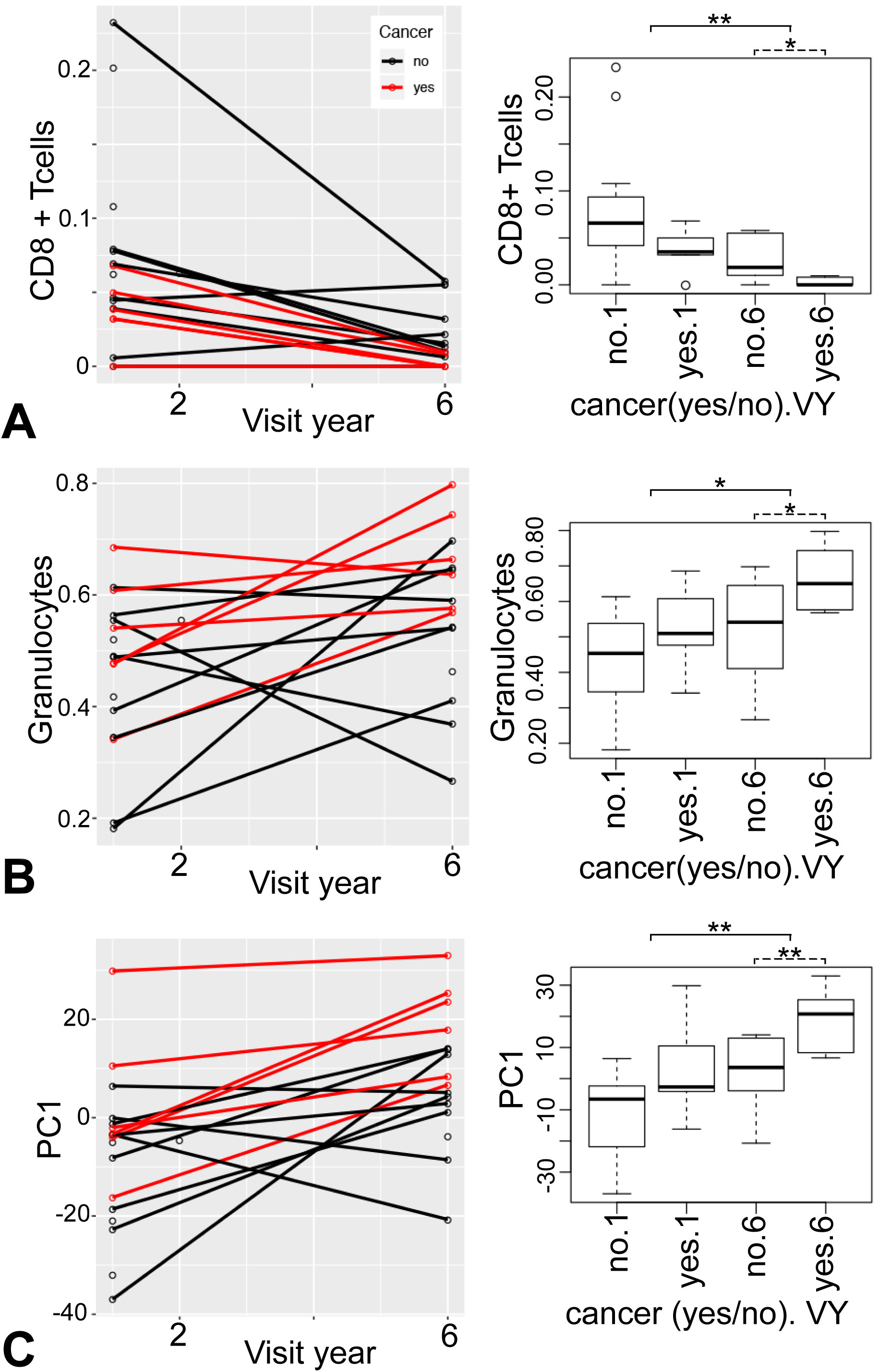
Longitudinal plots for DNA methylation-based estimates. The line plots (left) show the individual trajectory over time and the box plots (right) show the data averaged by visit year (baseline = 1, and Year 6) in cancer-free (no) or cancer-present (yes) groups. **(A)** Estimated proportions of CD8+ T-cells show a significant decline over time (baseline vs Year 6, solid line above boxplots) and are lower in the cancer-present group relative to the cancer-free group at Year 6 (cancer-free vs cancer-present, dashed line above boxplots). **(B)** Granulocyte proportions generally increase over time and are higher in the cancer-present group by Year 6. **(C)** The first principal component (PC1) computed from genome-wide methylation shows significant change over time as well as significant cross-sectional difference between the cancer-free and cancer-present groups by Year 6. In the line plots, red lines identify individuals who received a cancer diagnosis, and black lines identify those who remained cancer-free. Significance codes are *p-value < 0.05, **p-value < 0.01.

### Association between CpG-based blood cell estimates and cancer

We next examined if variation in blood cell composition was associated with cancer diagnosis. We performed the analysis stratified by baseline and Year 6. At baseline, none of the blood cells differentiated between those who developed cancer and those who remained cancer-free. By Year 6, CD8+ T-cell proportion was lower and granulocyte proportion was higher in the cancer-present group with modest statistical significance (**Figure 1A, B**; **Table 3**).

### Global patterns in DNA methylation and association with cell composition

To examine the global patterns of variation in the methylome, we performed PCA using the 739,648 probes. PC1 to PC5 captured 49% of the variance in the data (**Additional file 4: Data S1**). Age and BMI were not correlated with the top 5 PCs. PC4 showed an association with race only at Year 6 (p-value = 0.02), and PC5 with sex only at baseline (p-value = 0.02) (full results in **Additional file 4: Data S1**).

Correlation with blood cell estimates showed that PC1, which accounts for 21% of the variance, had a strong positive correlation with granulocytes and negative correlations with lymphoid cells (T-cells, B-cells, and natural killer or NK cells) at both baseline and Year 6 (full correlation matrix is provided in **Additional file 4: Data S1**). PC5 was positively correlated with monocytes at both baseline and Year 6 (**Additional file 4: Data S1**).

### Global patterns in DNA methylation and association with cancer

We next evaluated whether the PCs could differentiate between individuals who remained cancer-free compared to those who received a cancer diagnosis. PC1, which captured the variation in cellular composition, showed a modest association with cancer diagnosis at baseline and this became stronger by Year 6 (**Table 3**; **Figure 1C**). The remaining 4 PCs were not associated with cancer (**Additional file 4: Data S1**).

### Differential CpG methylation between cancer and cancer-free groups

Following the PC analysis, we explored differential methylation at the level of individual CpGs. Given the small sample size, we carried out simple t-tests to compare the cancer-present vs. cancer-free groups at Year 6, the time when PC1 showed a significant difference between the two groups. To control for cellular heterogeneity and unmeasured confounding variables, we performed the EWAS using residual β-values adjusted for the first 5 PCs. No CpG reached the genome-wide significant threshold (p-value ≤ 5 × 10^−8^). However, three CpGs, including one located in an intronic CpG island of the metastasis associated gene (cg02162462, *MTA3*), were genome-wide suggestive (p-value ≤ 10^−5^) (**Figure 2**). We considered the top 10 cancer-associated CpGs and evaluated these for replication (**Table 4**). Among these top 10, 5 CpGs were associated with lower methylation in the cancer group (cancer-hypomethylated), and the remaining 5 showed higher methylation in the cancer group (cancer-hypermethylated). To test for replication, we cross-checked our results with those from Roos et al., which evaluated for pan-cancer CpG biomarkers in blood using the previous version of the Illumina Human Methylation 450K (HM450K) array. [16]. Of the top 10 CpGs in **Table 4**, 5 probes were also represented in the HM450K array. The CpG in the intron of *RPTOR* (cg08129331), which was cancer-hypomethylated in Health ABC, also showed a similar hypomethylation in the Roos cohort at p-value = 0.05. The CpG in the 3’ UTR of *MRPL44*, which showed cancer-hypermethylation in Health ABC, showed hypermethylation in the Roos cohort at p-value = 0.08.

**Table 4.** Top 10 cancer associated CpGs.

| ProbeID | Chr (Mb) | Location<sup>2</sup> | Residual β-value<br>Y6 ttest in<br>HABC<sup>3</sup> | Replication<br>in Roos. et<br>al. | Correlation<br>of Y6-Y1<br>with YTD in<br>HABC<sup>5</sup> |
| --- | --- | --- | --- | --- | --- |
|  |  |  | Canc. yes-<br>no<br>(pval) | Canc. yes-<br>no<br>(pval)<sup>4</sup> | R |
| cg09608390 | 17(1.00) | exon *ABR* | 0.019<br>(1.1E-06) |  | -0.42<br>(0.41) |
| cg01399430 | 5(6.52) | intergenic | -0.048<br>(5.6E-06) |  | 0.34<br>(0.50) |
| cg02162462 | 2(42.8) | Intron1 *MTA3*;<br>CGI | -0.027<br>(1.0E-05) | 0.02<br>(0.93) | 0.63<br>(0.18) |
| cg25105842 | 2(224.83) | 3'UTR *MRPL44* | 0.016<br>(1.6E-05) | 0.37<br>(0.08) | 0.09<br>(0.86) |
| cg05808305 | 11(2.77) | intron; *KCNQ1* | -0.016<br>(1.8E-05) |  | 0.30<br>(0.57) |
| cg25403416 | 19(30.19) | 3'UTR;<br>*C19orf12* | 0.019<br>(1.8E-05) | -0.14<br>(0.30) | -0.06<br>(0.91) |
| cg07516252 | 14(24.64) | promoter<br>*REC8*; CGI | -0.038<br>(2.0E-05) | 0.08<br>(0.37) | **0.89**<br>**(0.02)** |
| cg08129331 | 17(78.56) | Intron1 *RPTOR* | -0.039<br>(2.4E-05) | **-0.13**<br>**(0.05)** | **0.83**<br>**(0.04)** |
| cg11784099 | 21(46.23) | Intron1 *SUMO3* | 0.035<br>(2.4E-05) |  | -0.06<br>(0.91) |
| cg04429789 | 1(45.52) | intron *ZSWIM5* | 0.024<br>(2.7E-05) |  | **-0.81**<br>**(0.05)** |
| CD8T |  |  | -0.027<br>(0.02) |  | -0.202<br>(0.70) |
| Gran |  |  | 0.14<br>(0.04) |  | -0.05<br>(0.93) |
| PC1 |  |  | 17.01(0.008) |  | -0.21<br>(0.69) |

<sup>1</sup> GRCh37/hg19
<sup>2</sup> CGI is CpG island
<sup>3</sup> Mean difference between cancer-present and cancer-free groups of Health ABC at Year 6 and t-test p-values
<sup>4</sup> Mean difference between cancer discordant twins in Roos et al. (yes – no) and t-test p-values
<sup>5</sup> Mean Correlation between years to diagnosis and longitudinal change in residual β-values (deltaβ = Year 6 – baseline) in cancer group of Health ABC

**Figure 2:**
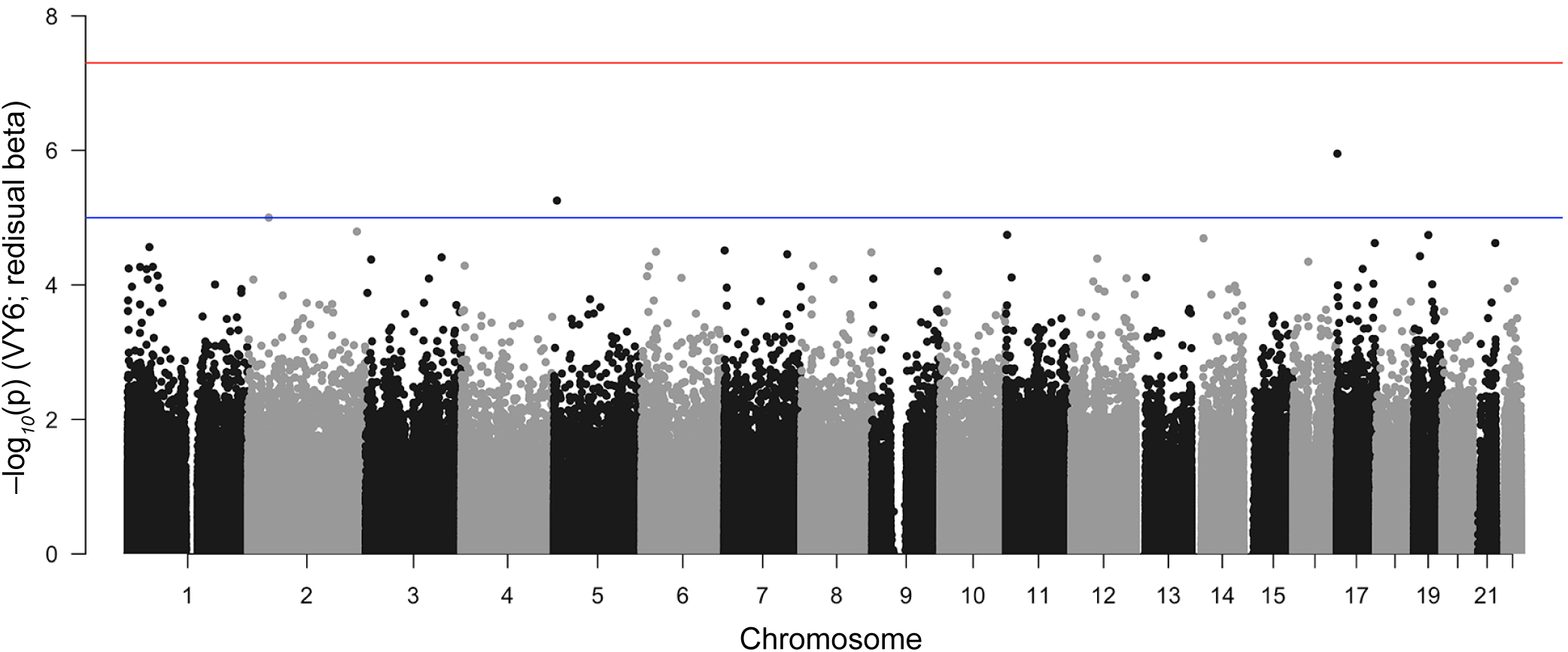
Epigenome-wide association plot. The Manhattan plot shows the association between the CpGs and cancer at Year 6. The x-axis represents the chromosomal locations, and each point depicts a CpG probe. The y-axis is the –log_10_(p-value) of differential methylation between those who received cancer diagnosis vs. those who remained cancer-free. The red horizontal line indicates the genome-wide significant threshold (p-value ≤ 5 × 10^−8^) and the blue horizontal line indicates the suggestive threshold (p-value ≤ 10^−5^).

### Longitudinal changes in CpG methylation and diagnosis time

Since these CpGs differentiated between those who developed cancer and those who remained cancer-free at Year 6, we then explored if the longitudinal changes in methylation over time (deltaβ = Year 6 – baseline) could be related to time to cancer diagnosis. For the 5 cancer-hypomethylated CpGs in **Table 4**, we predicted that the within-individual decline in methylation at Year 6 (negative deltaβ) would be greater in those who were closer to diagnosis (positive correlation with years to diagnosis or YTD). Inversely, for the 5 cancer-hypermethylated CpGs, we predicted that the within-individual increase in methylation at Year 6 (positive deltaβ) would be greater in those closer to diagnosis (negative correlation with YTD). With the exception of three probes that showed Pearson correlation near 0, the remaining seven CpGs showed a correlation pattern that was consistent with our predictions (**Table 4**). The CpGs in *REC8* (cg07516252), *RPTOR*, and *ZSWIMS* (cg04429789) were statistically significant at p-value ≤ 0.05. **Figure 3** shows the longitudinal plots for these 3 CpGs and the correlation between deltaβ and YTD.

**Figure 3:**
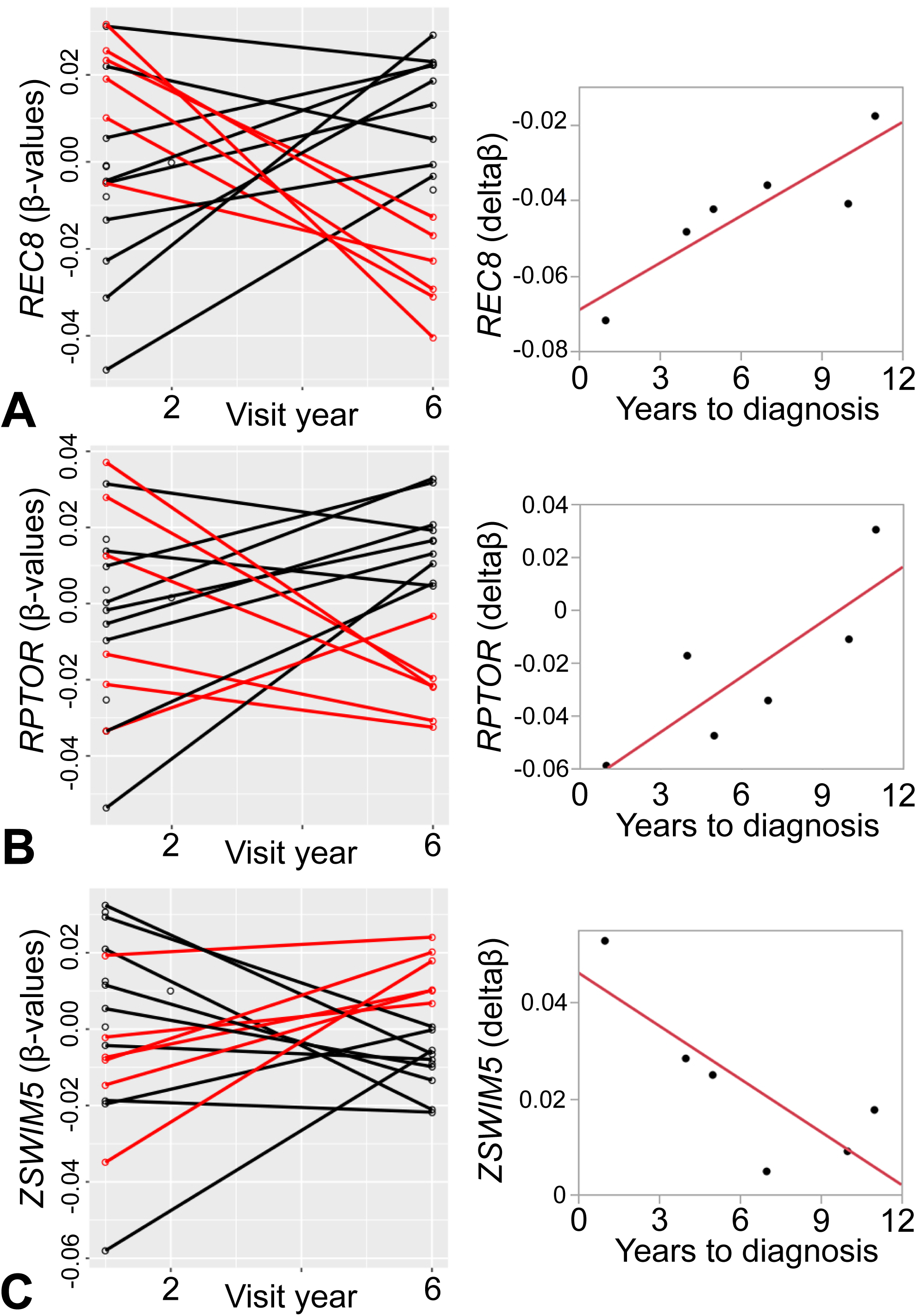
Longitudinal rate of change in CpG methylation. The line plots (left) show the individual DNA methylation β-values from baseline to Year 6 for CpGs in **(A)** *REC8* (cg07516252), **(B)** *RPTOR* (cg08129331), and **(C)** *ZSWIM5* (cg04429789). Red lines identify individuals who received a cancer diagnosis, and black lines identify those who remained cancer-free. Longitudinal changes in DNA methylation were calculated as deltaβ =Year 6 – baseline, and the correlations between deltaβ and years to cancer diagnosis are shown for the respective CpGs (right). Higher magnitude of change is seen in individuals closer to clinical diagnosis.

## Discussion

### Summary

In this study, we evaluated two aspects of the aging methylome in an older group of participants: (1) differences in DNA methylation patterns between those who developed cancer and those who remained cancer-free, and (2) the longitudinal trajectory over time. We used DNA purified from peripheral blood cells collected from a subset of Health ABC Study participants who provided DNA samples separated by approximately 5 years. Overall, there was strong intra-individual stability from baseline to Year 6, and with the exception of two participants, all other participants with longitudinal samples paired with self when grouped by unsupervised hierarchical clustering. When a large number of random CpGs or genome-wide data are used in such clustering analysis, samples generally group by age and shared genotype (i.e., either monozygotic twins or with self), with few exceptions [26–28]. The few exceptions likely reflect individual discordance and epigenetic drift that occurs within a person, particularly at old age [24, 25]. We found that cellular composition is a major source of variation and significantly contributed to the variance explained by the primary principal component (PC1). In terms of the biomarker utility of DNA methylation, our study highlighted a few CpGs as potential biomarkers, and the dynamic changes over time at these CpGs were correlated with time to cancer diagnosis.

### Cellular heterogeneity as both informative and a potential confounder

Cellular composition is clearly a major correlate of DNA methylation and can be a confounding variable when we attempt to relate the methylome derived from heterogeneous tissue to aging and disease [29]. The composition of cells in circulating blood can be influenced by natural immune aging and also by numerous correlated health variables including lifestyle, infectious disease, leukemia or similar cancers, and environmental exposures. For example, one of the most consistent features of the aging immune system involves thymic involution and the time-dependent decline in both the absolute number and the relative percent of naïve CD8+ T-cells [30–33]. A strategy to estimate the composition of cells from DNA methylation data is to rely on specific CpGs that are known to be strong cell-specific markers and can serve as surrogate measures of cellular sub-types [21–23]. With the current data, we applied this *in silico* approach to estimate the relative proportions of CD8+ T-cells, CD4+ T-cells, B-cells, NK cells, granulocytes, and monocytes. The DNA methylation-based estimates of cell proportions showed a decrease in CD8+ T-cells and an increase in granulocytes over the course of 5 years. By Year 6 from baseline, the proportion of CD8+ T-cells was lower and proportion of granulocytes higher in the cancer-present group relative to the cancer-free group. Since the first few PCs captured the variance due to cellular composition, PC1 also showed a similar change over time. PC1 showed a slight distinction between the cancer-present vs. cancer-free groups even at baseline, and this became more pronounced by Year 6. These differences are likely because PC1 summarized the changes in the composition of multiple cell subtypes including those that were not estimated using the reference set of cell-specific CpGs. PCA may therefore be more effective at capturing the composite changes arising from different cellular subtypes and may also be more disease-informative than the estimated proportion of major cell types.

Our observations are consistent with the general decrease in lymphoid cells and increase in myeloid cells during aging [30–32]. In line with the lower lymphocytes and higher granulocytes in the cancer group, work from both model organisms and humans have shown an inverse relationship between lymphocytes and granulocytes with lower B-cells and T-cells, and higher neutrophils being associated with higher mortality risk [34–36]. While we cannot disentangle the inter-correlations between aging, cell composition, and methylation patterns, our results do demonstrate that DNA methylation data derived from peripheral blood in older participants can be used to glean information on their cellular profiles, and this in turn can be related to their health and disease status.

### Identifying (pan)cancer CpGs

Following the cell estimation and PC analysis, we took an EWAS approach to examine differential methylation at the level of individual CpGs. Previous studies have already demonstrated that DNA methylation patterns can provide a powerful “pan-cancer” biomarker—i.e., an epigenetic signature of cancer that can serve as a general biomarker for the presence of cancer, and possibly different cancer types as well [37, 38]. The majority of these studies have involved comparisons between normal vs. tumor tissue, or are dependent on the shedding of cell-free DNA from the primary site of cancer and therefore are indicators of *in situ* changes that occur in tumor cells [37, 39-43]. Relatively few studies have taken a prospective approach that involves sample collection prior to disease diagnosis [44, 45], and even fewer have attempted to track longitudinal changes across multiple timepoints [14, 15]. Nevertheless, these few prospective studies have shown that both the global patterns and DNA methylation at specific CpG sites can be indicators of cancer, and even more strikingly, that some of these generalized changes can be detected in circulating blood cells [14, 15, 44, 45].

Given this background, our goal was to examine if we can also detect similar “pan-cancer” CpG biomarkers. We used a simple approach and contrasted DNA methylation between the cancer-present and cancer-free groups at Year 6, the time when we expect the differences to be more pronounced. Despite the small sample size, 3 CpGs passed the conventional genome-wide suggestive threshold of 10^−5^ [46], and the suggestive hits included a CpG located in the first intron and overlapping a CpG island within the metastasis associated 1 family member 3 (*MTA3*), a gene known to play a role in tumorigenesis and metastasis. To incorporate the longitudinal information, we then focused on the top 10 differentially methylated CpGs and examined whether the within-individual longitudinal changes in β-values in the cancer group were correlated with time to diagnosis. Due to the small sample size, it was not feasible to evaluate correlations with cancer stage or progression, and the correlations were examined only for the time to the first adjudicated diagnosis. The overall trend indicated that the magnitude of change over five years, with greater negative slope for cancer-hypomethylated CpGs and correspondingly greater positive slope for cancer-hypermethylated CpGs, was correlated with the time to cancer diagnosis. Although this analysis was carried out in only the 6 cancer cases, the correlations between deltaβ and time to diagnosis were significant for the CpGs in the promoter region of *REC8*, and introns of *RPTOR* and *ZSWIM5*.

To gather additional lines of evidence, we examined if the association with cancer for these CpGs can be replicated in an independent dataset, and if the cognate genes have been previously related to cancer or tumorigenesis. For replication we referred to the work by Roos et al. [16]. While the study by Roos et al. compared cancer-discordant monozygotic twins and involved a much wider age range, some design features common to our study are: (1) the cancer group included samples collected from individuals who had already received cancer diagnosis (post-diagnosis) and from individuals within 5 years to diagnosis (pre-diagnosis), (2) a variety of cancer types were represented, and (3) genome-wide DNA methylation was measured using peripheral blood cells. In the Health ABC Study set, 3 participants (excluding Per13 with leukemia) had been diagnosed by Year 6, and the remaining participants received a diagnosis 1–5 years after Year 6. Since the Roos dataset was generated on the previous version of the Illumina DNA methylation arrays (HM450K), only 5 of the top 10 probes were represented on that array and could be evaluated for replication. Only the CpG in the intron of *RPTOR* (cg08129331) was replicated and was also associated with a consistently lower methylation in the cancer group (p-value = 0.05 in Roos study). The 3’UTR CpG in *MRPL44* (cg25105842) showed a consistent increase in methylation in the Roos study, but this did not reach statistical significance (p-value = 0.08).

### Cancer associated CpGs in tumor suppressor genes

Eight of the top ten cancer CpGs were located within annotated gene features including the top CpG, cg09608390, located in the exon of RhoGEF and GTPase activating protein gene, *ABR*. We did not find a clear-cut link between *ABR* and cancer in the existing literature. However, among the eight genes in the list, *REC8* (meiotic recombination protein) is a known tumor suppressor. There is also evidence that *KCNQ1* (potassium voltage-gated channel member), *MTA3*, and *ZSWIM5* (zinc finger SWIM-type 5) have tumor suppressive roles.

*MTA3* is a chromatin remodeling protein that has a complex association with cancer [47, 48]. In certain types of malignant tumors such as glioma, certain breast cancers, and adenocarcinomas, *MTA3* is under-expressed and is implicated as a tumor suppressor [48–51]. In other carcinomas such as hepatocellular, lung, gastric, and colorectal cancers, *MTA3* is reported to be overexpressed, with higher expression correlated with tumor progression and poorer prognosis [52–56]. In the Health ABC samples, the CpG (cg02162462) located in the first intron of *MTA3* and overlapping a CpG island had lower methylation in the cancer-present group at Year 6. At baseline, there was no significant difference between the groups. The negative deltaβ, though not statistically significant, was greater in participants closer to receiving a clinical cancer diagnosis (Pearson correlation R = 0.63). While we could not replicate this CpG in the Roos dataset, the collective evidence suggests that methylation changes in the CpG island of *MTA3* may be associated with tumor development and progression.

*REC8* has a more consistent tumor suppressive role and promoter hypermethylation and suppression of its expression occurs in tumor cells [57–60]. In the Health ABC samples, the CpG in the promoter (cg07516252) was hypomethylated and not hypermethylated in the group that received cancer diagnosis. The rate of promoter hypomethylation was also significantly correlated with time to diagnosis (R = 0.89). Since our study is blood-based and does not stem from the primary tumor site, the hypomethylation may indicate aberrant methylation over time in individuals, with greater changes observed in those individuals who are closer to clinical manifestations. However, this promoter CpG did not replicate in the Roos data.

*KCNQ1* is another tumor suppressor gene, and loss of its expression is considered to be an indicator of metastasis and poor prognosis [61–63]. There is also evidence that the reduction in *KCNQ1* expression in cancer cells may be mediated by promoter hypermethylation [62, 64]. In the Health ABC samples, the intronic CpG (cg05808305) had much lower methylation in the cancer group and was significant only at Year 6. Among the known and potential tumor suppressive genes, only the intronic CpG in *ZSWIM5* (cg04429789) was associated with hypermethylation in the Health ABC cancer diagnosed group; for this CpG, the positive deltaβ was significantly correlated with time to diagnosis with greater positive change in those closer to receiving a diagnosis (R = −0.81). So far, we have found only one study showing that the expression of *ZSWIM5* inhibits malignant progression [65]. We could not test replication for the CpG in *ZSWIM5* since this was not a probe that was included in the HM450K array.

Based on the multiple lines of evidence, we highlight the CpG in the first intron of *RPTOR* (cg08129331) as a stronger potential pan-cancer biomarker as this specific CpG was replicated in the Roos data. This gene codes for a member of the mTOR protein complex, which plays a key role in cell growth and proliferation, and dysregulation of this signaling pathway is a common feature in cancers [66]. The lower methylation of this CpG in cancer-free individuals in Health ABC was significant only in Year 6. For the longitudinal change, the correlation between the deltaβ and time to diagnosis was significant for cg08129331. This specific CpG has been previously presented as a marker to differentiate between different medulloblastoma subtypes [67]. Another study has also indicated that the decrease in methylation in *RPTOR* measured in peripheral blood may be a biomarker for breast cancer, although this failed replication in a follow-up study [68, 69]. Similar to *REC8*, there was more negative change in β-value from Year 1 to 6 in individuals closer to receiving a cancer diagnosis.

### Limitations

The present work was carried out in a very small and heterogenous group of participants. The cancer-present group consisted of different types of cancers, and there was a combination of individuals who received the diagnosis before and after Year 6. The differences in DNA methylation should therefore be interpreted as potential correlates rather than predictive indicators of disease. Due to the limitation in sample number, we performed simple t-test comparisons rather than more complex regressions such as mixed modeling. Furthermore, we considered the cancer diagnosis as the main outcome variable and did not account for cancer type, stage or progression. Additionally, while we took steps to statistically correct for immune cell composition, the data was derived from white blood cells from older participants. The *in-silico* approach to estimate cell composition cannot discern the finer repertoire of cellular subtypes that are known to change particularly in older individuals. The results we present therefore require further replication in a larger cohort. Our study is mainly a demonstration of concept that highlights the utility of longitudinal blood collection and the potential information on health and disease that can be gained by tracking dynamic changes in the methylome.

### Conclusion

Taken together, our analysis detected global changes in the methylome that are partly due to cellular heterogeneity and also due to changes at specific CpGs that could indicate cancer development and progression. From the multiple lines of evidence, we posit methylation in *RPTOR* as a potential biomarker of cancer that justifies further investigation and validation.

## Supporting information

Additional file 1: Figure S1. Microarray data quality checks (A) The density plots for ??-values using the full set of 866,836 probes show the expected

Additional file 2: Figures S2. Samples pair by participant ID. Unsupervised hierarchical clustering using probes that were flagged due to overlap with

Additional file 3: Table S1. DNA methylation-based estimation of blood cell proportions

Additional file 4: Data S1. Analysis of top 5 principal components and association with demographics, blood cell estimates, and cancer diagnosis

## Abbreviations

CGI: CpG island
EA: European Americans or Caucasians
AA: African Americans
EWAS: epigenome-wide association studies
Gran: granulocytes
Health ABC: Health, Aging and Body Composition Study
HM850K: Illumina Infinium Human MethylationEPIC
HM450K: Illumina Human Methylation 450K
NK cells: natural killer cells
PCA: principal component analysis
PC: principal components
QC: quality checks
SNP: single nucleotide polymorphism
YTD: years to diagnosis

## Declarations

### Ethics approval and consent to participate

All participants provided written informed consent and all Health ABC Study sites received IRB approval

### Consent for publication

Not applicable

### Data availability

Full raw data normalized data and full EWAS statistics will be deposited to the NCBI NIH Gene Expression Omnibus (will be submitted upon acceptance by a peer-reviewed journal).

### Author contributions

AHB: performed analysis and contributed to the writing of the manuscript; JWL and JVSL: helped with data analysis and contributed to the final manuscript; JHF and KCJ: helped with interpretation and contributed to the writing of the manuscript; EMS: helped with interpretation and contributed to the final manuscript; KM designed the study, contributed to data analysis and interpretation, and prepared the manuscript.

## Acknowledgements

We are very thankful to the Health ABC Study for granting us access to the data and DNA samples. We also express our deep gratitude to the late Dr. Suzanne (Suzy) Satterfield, M.D., for the help and advice she provided. This work was supported by funds from UTHSC Faculty Award UTCOM-2013KM. The Health ABC Study research was funded in part by the Intramural Research Program of the NIH, National Institute on Aging (NIA), and supported by NIA Contracts N01-AG-6-2101; N01-AG-6-2103; N01-AG-6-2106; NIA grant R01-AG028050, and NINR grant R01-NR012459.

## List and description of additional files

**Additional file 1: Figure S1. Microarray data quality checks**

**(A)** The density plots for β-values using the full set of 866,836 probes show the expected bimodal distribution. **(B)** Unsupervised hierarchical clustering using the full set of probes shows that, with the exception of two participants (Per1 and Per9), all samples with longitudinal data pair appropriately with self. This cluster tree identifies Per13 as an outlier at both baseline and visit year 6. **(C)** Principal component analysis was done using a filtered set of 739,648 autosomal probes. The scatter plot between principal component 1 (PC1) and PC2 identifies Per13 as an outlier.

**Additional file 2: Figures S2. Samples pair by participant ID.**

Unsupervised hierarchical clustering using probes that were flagged due to overlap with SNPs shows that samples collected longitudinally from the same participant pair perfectly.

**Additional file 3: Table S1. DNA methylation-based estimation of blood cell proportions**

**Additional file 4: Data S1. Analysis of top 5 principal components and association with demographics, blood cell estimates, and cancer diagnosis**

