## Supplementary figures and images for "Longitudinal Study of Leukocyte DNA Methylation and Biomarkers for Cancer Risk in Older Adults"

### Additional file 1: Figure S1. Microarray data quality checks (A) The density plots for ??-values using the full set of 866,836 probes show the expected

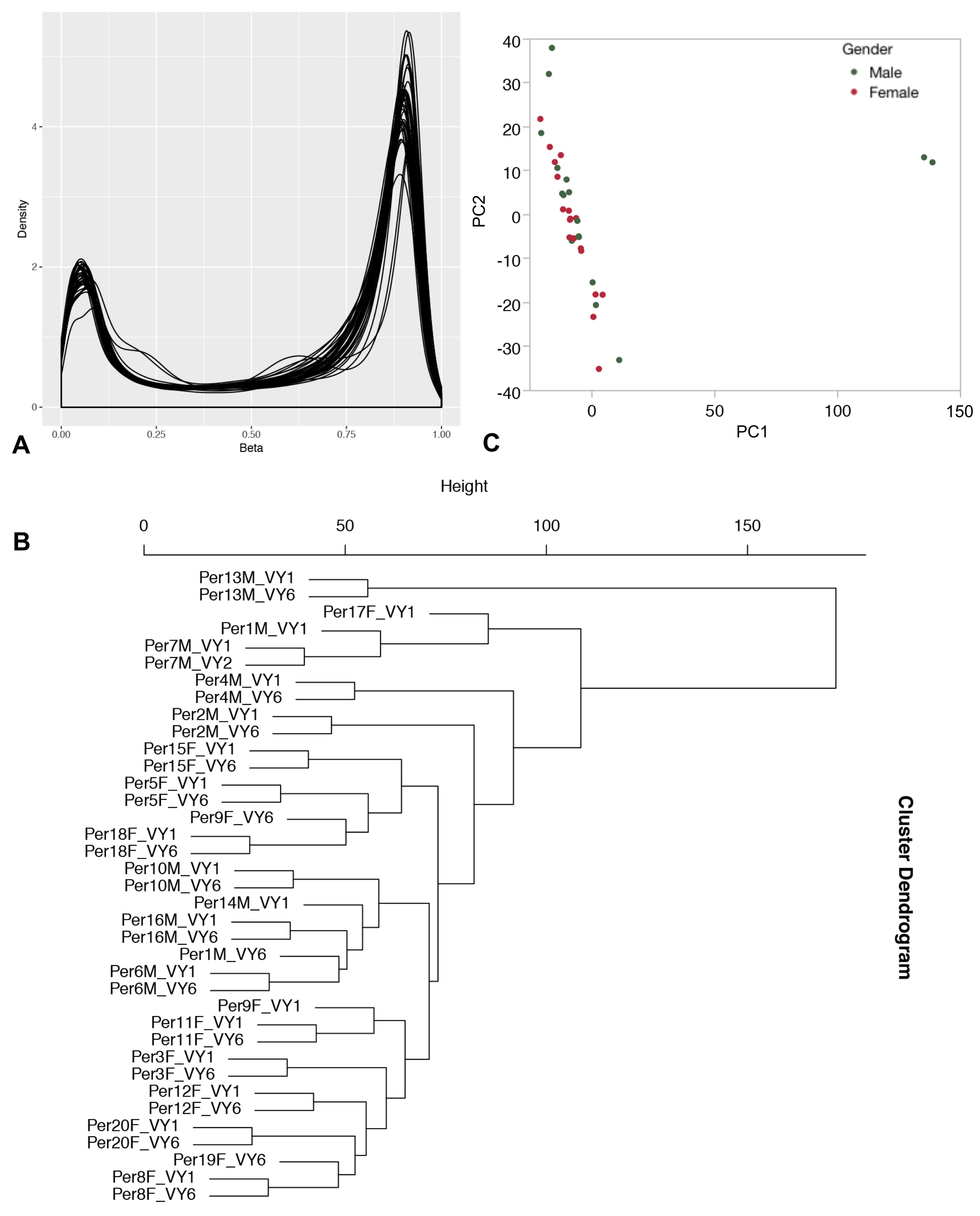

### Additional file 2: Figures S2. Samples pair by participant ID. Unsupervised hierarchical clustering using probes that were flagged due to overlap with

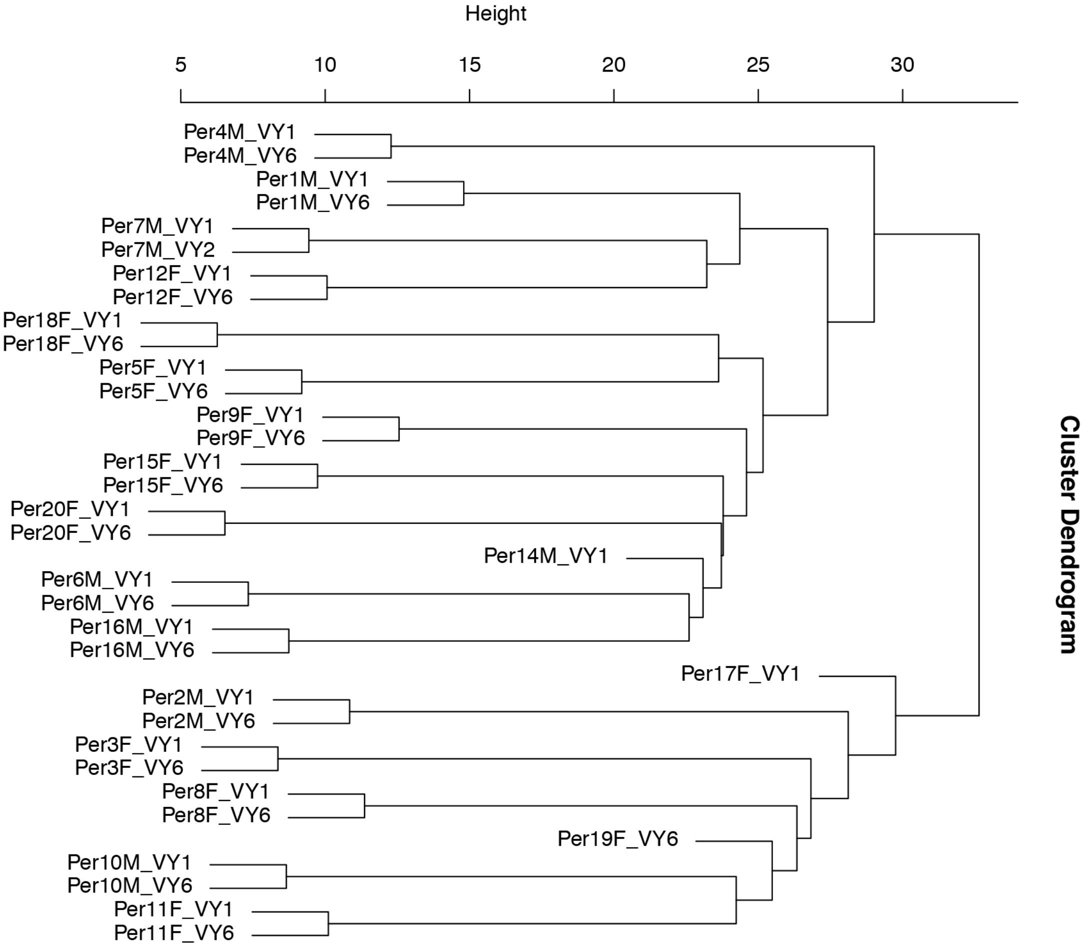
