## Additional file 3: Table S1. DNA methylation-based estimation of blood cell proportions for "Longitudinal Study of Leukocyte DNA Methylation and Biomarkers for Cancer Risk in Older Adults"

SUPPLEMENTAL TABLE

**Table S1. DNA methylation-based estimation of blood cell proportions**

| **ID** | **Canc.** | **Visit** | **CD8T** | **CD4T** | **NK** | **Bcell** | **Mono** | **Gran** | **Age** |
| --- | --- | --- | --- | --- | --- | --- | --- | --- | --- |
| Per1 | no | Baseline | 0.23 | 0.16 | 0.23 | 0.08 | 0.16 | 0.18 | 75 |
|  |  | Year 6 | 0.06 | 0.07 | 0.08 | 0.03 | 0.07 | 0.70 | 81 |
| Per2 | yes | Baseline | 0.07 | 0.27 | 0.08 | 0.06 | 0.07 | 0.48 | 71 |
|  |  | Year 6 | 0.01 | 0.09 | 0.03 | 0.04 | 0.05 | 0.80 | 77 |
| Per3 | no | Baseline | 0.07 | 0.21 | 0.20 | 0.13 | 0.24 | 0.19 | 72 |
|  |  | Year 6 | 0.03 | 0.16 | 0.13 | 0.06 | 0.24 | 0.41 | 78 |
| Per4 | yes | Baseline | 0.00 | 0.19 | 0.03 | 0.01 | 0.09 | 0.69 | 74 |
|  |  | Year 6 | 0.00 | 0.18 | 0.05 | 0.09 | 0.12 | 0.64 | 80 |
| Per5 | no | Baseline | 0.08 | 0.09 | 0.16 | 0.04 | 0.09 | 0.56 | 76 |
|  |  | Year 6 | 0.01 | 0.10 | 0.12 | 0.02 | 0.11 | 0.65 | 82 |
| Per6 | yes | Baseline | 0.03 | 0.15 | 0.17 | 0.06 | 0.07 | 0.54 | 75 |
|  |  | Year 6 | 0.00 | 0.08 | 0.16 | 0.08 | 0.12 | 0.58 | 81 |
| Per7 | no | Baseline | 0.11 | 0.17 | 0.19 | 0.07 | 0.10 | 0.42 | 76 |
|  |  | Year 2 | 0.06 | 0.13 | 0.16 | 0.03 | 0.09 | 0.56 | 78 |
| Per8 | no | Baseline | 0.04 | 0.21 | 0.11 | 0.09 | 0.08 | 0.49 | 78 |
|  |  | Year 6 | 0.01 | 0.20 | 0.09 | 0.10 | 0.08 | 0.54 | 84 |
| Per9 | yes | Baseline | 0.04 | 0.19 | 0.15 | 0.09 | 0.11 | 0.48 | 78 |
|  |  | Year 6 | 0.00 | 0.09 | 0.06 | 0.03 | 0.08 | 0.74 | 84 |
| Per10 | yes | Baseline | 0.03 | 0.29 | 0.19 | 0.05 | 0.12 | 0.34 | 74 |
|  |  | Year 6 | 0.00 | 0.21 | 0.10 | 0.03 | 0.11 | 0.57 | 80 |
| Per11 | no | Baseline | 0.05 | 0.32 | 0.18 | 0.08 | 0.08 | 0.34 | 74 |
|  |  | Year 6 | 0.02 | 0.23 | 0.09 | 0.04 | 0.11 | 0.54 | 80 |
| Per12 | no | Baseline | 0.04 | 0.14 | 0.12 | 0.09 | 0.08 | 0.56 | 71 |
|  |  | Year 6 | 0.06 | 0.20 | 0.28 | 0.06 | 0.16 | 0.27 | 77 |
| Per13 | yes | Baseline | 0.00 | 0.16 | 0.04 | 0.64 | 0.08 | 0.00 | 76 |
|  |  | Year 6 | 0.00 | 0.17 | 0.05 | 0.65 | 0.06 | 0.00 | 82 |
| Per14 | no | Baseline | 0.06 | 0.20 | 0.09 | 0.08 | 0.07 | 0.52 | 75 |
| Per15 | no | Baseline | 0.08 | 0.29 | 0.09 | 0.06 | 0.10 | 0.39 | 73 |
|  |  | Year 6 | 0.01 | 0.18 | 0.07 | 0.04 | 0.07 | 0.65 | 79 |
| Per16 | no | Baseline | 0.00 | 0.18 | 0.10 | 0.04 | 0.10 | 0.61 | 73 |
|  |  | Year 6 | 0.00 | 0.19 | 0.09 | 0.08 | 0.10 | 0.59 | 79 |
| Per17 | no | Baseline | 0.20 | 0.20 | 0.08 | 0.10 | 0.12 | 0.35 | 76 |
| Per18 | yes | Baseline | 0.05 | 0.14 | 0.09 | 0.02 | 0.11 | 0.61 | 78 |
|  |  | Year 6 | 0.01 | 0.11 | 0.10 | 0.01 | 0.12 | 0.66 | 84 |
| Per19 | no | Year 6 | 0.06 | 0.31 | 0.06 | 0.04 | 0.09 | 0.46 | 78 |
| Per20 | no | Baseline | 0.01 | 0.22 | 0.17 | 0.04 | 0.09 | 0.49 | 70 |
|  |  | Year 6 | 0.02 | 0.29 | 0.17 | 0.03 | 0.13 | 0.37 | 76 |
